# A combined strategy of neuropeptide predictions and tandem mass spectrometry identifies evolutionarily conserved ancient neuropeptides in the sea anemone *Nematostella vectensis*

**DOI:** 10.1101/593384

**Authors:** Eisuke Hayakawa, Hiroshi Watanabe, Gerben Menschaert, Thomas W. Holstein, Geert Baggerman, Liliane Schoofs

## Abstract

Neuropeptides are a class of bioactive peptides and are responsible for various physiological processes including metabolism, development and reproduction. Although accumulated genome and transcriptome data have reported a number of neuropeptide candidates, it still remains difficult to obtain a comprehensive view of neuropeptide repertoires due to their small and variable nature. Neuropeptide prediction tools usually work only for those peptides for which sequentially related homologs have previously been identified. Recent peptidomics technology has enabled systematic structural identification of neuropeptides by using the combination of liquid chromatography and tandem mass spectrometry. However, obtaining reliable identifications of endogenous peptides is still difficult using a conventional tandem mass spectrometry-based peptide identification approach using protein database because a large search space has to be scanned due to the absence of a cleavage enzyme specification. We developed a pipeline consisting of the prediction of *in silico* cleaved endogenous neuropeptides followed by peptide-spectrum matching enabling highly sensitive and reliable neuropeptide identification. This approach effectively reduces the search space of peptide-spectrum matching, and thus increases search sensitivity. To identify neuropeptides in *Nematostella vectensis,* a basal eumetazoan having one of the most primitive nervous systems, we scanned the *Nematostella* protein database for sequences displaying structural hallmarks of metazoan neuropeptides, including C/N-terminal structures and modifications. Peptide-spectrum matching was performed against the *in silico* cleaved peptides and successfully identified dozens of neuropeptides at high confidence. The identification of Nematostella neuropeptides structurally related the tachykinin, GnRH/AKH, neuromedin-U/pyrokinin peptide families indicate that these peptides already originated in the eumetazoan ancestor of all animal species, most likely concomitantly with the development of a nervous system.

## Introduction

Neuropeptides are a highly diverged group of messenger molecules and function as essential communicators for fairly standard physiological processes such as muscle contraction, food digestion, growth, development, reproduction as well as for more complex actions and long-term effects such as behavioural adaptation, memory formation and even ageing. A neuropeptide is usually encoded in a larger neuropeptide precursor gene that also typically encodes an N-terminal signal peptide. The precursor is translated on the rough endoplasmic reticulum, and the signal peptide is removed by a signal peptidase. Afterwards, processing enzymes in the immature secretory granules typically produce a mature neuropeptide by processing the precursor at cleavage sites. A number of cleavage enzymes with specific recognition patterns have been currently identified [1,2]. After cleavage, the cleaved peptides can undergo various post translational modifications (PTMs) [3]. The N-terminal or C-terminal residues are often modified and these modifications often contribute to the stability of the mature peptides. The most frequently observed modification is amidation at the C-terminus. It occurs in many neuropeptides and is often required for their biological activities. Enzymatic transformation of the C-terminus from a glycine to an alpha-amide is catalyzed by peptidylglycine alpha-amidating monooxygenase [4]. First, peptidylglycine is transformed into peptidyl-alpha-hydroxyglycine in the presence of copper, ascorbate, and molecular oxygen and then converted to peptide alpha-amide and glyoxylate. C-terminal amidation protects the mature peptide from enzymatic degradation from C-terminus. Pyroglutamic acid is widely observed at N-termini of various neuropeptides and protects the peptide chain from enzymatic degradation from N-terminus. In addition to PTMs, many bioactive peptides contain a proline residue in the second or third position from the N-terminus. This feature also protects the peptide from peptidase activity at the N-terminus [5]. All these structural characteristics can be widely observed in neuropeptide sequences across the Animal Kingdom [6].

Recent advances in mass spectrometry and liquid chromatography technology have led to the establishment of peptidomics, an efficient tandem mass spectrometry-based approach to identify neuropeptides. The combination of tandem mass spectrometry and peptide-spectrum matching tools enables systematic neuropeptide identification [7–9]. Unlike neuropeptide prediction tools that are based on sequence similarities, this peptidomics approach allows providing evidence for the *in vivo* presence of the cleaved peptides, in addition to determining the presence of PTMs of the peptide sequences [10,11]. However, in search of endogenous small peptides using conventional peptide-spectrum matching tools, achieving reliable identifications still remains difficult. Unlike proteomics, in which *in vitro* generated enzymatic digests are identified, cleavage sites in a precursor of endogenous neuropeptide cannot readily be accurately predicted. Therefore peptide-spectrum matching has to be performed without any enzyme specification, leading to a huge search space often resulting in poor identification confidence values. Besides, peptide-spectrum matching has to be done taking PTMs into consideration, even further increasing the search space [12].

Another and rather commonly employed approach to search for neuropeptides are homology-based database searches. Recent strides in sequencing genome and transcriptome in a wide variety of animal species have predicted bioactive peptides *in silico* [13–16]. Several researches successfully track down the evolutionarily conserved neuropeptides within a phylum or between closely related phyla, by means of sequence homology searching against protein sequence datasets [17]. This approach is very useful for searching peptide sequences that have been evolutionarily conserved between closely related species.. However, when peptides have become evolutionarily diverged, such as is the case for evolutionarily ancient organisms, the homology-based prediction of new neuropeptides may not always be successful. On top of this difficulty, within a particular peptide sequence, only a short motif required for the peptide’s biological activity, has in general been conserved during evolution [16,18] whereas the sequence of the none-peptide coding region of the precursor, is much less or not at all conserved. All these issues hamper reliable sequence similarity-based peptide prediction. Especially for evolutionarily distant species, it remains difficult to identify neuropeptides and corresponding genes based on homology searches and experimental validation is required in these cases. To deal with these hurdles, we have developed an alternative strategy to identify neuropeptides with high sensitivity and confidence. The strategy is a combination of the reduction of protein datasets to potential mature neuropeptide structures predicted based on the typical hall marks(signal peptide, specific cleavage sites, PTMs), combined with peptide-spectrum matching. The first step is effective in narrowing the search space for the peptide-spectrum matching and as such enables high sensitive peptide identifications. Based on this strategy, we here present a systematic identification of neuropeptides from *Nematostella vectensis*, one of the most evolutionarily ancient animals with a nervous system. We successfully identified 20 neuropeptides, many of which have not been predicted or annotated as neuropeptides when the genome of this organism was sequenced [19]. Compared to a conventional peptidomics workflow, this approach enables the identification of neuropeptides with the highest sensitivity.

## Materials and Methods

### Sample preparation

Adult polyps of *Nematostella vectensis* were kept in 1/3 artificial seawater at 18 °C and fed three time per week with nauplius larvae of *Artemia salina*. The culture medium was changed once a week. Five adult polyps were homogenized with a probe sonicator in 1.5 ml methanol/water/formic acid (FA) solution (90:9:1) in 3 cycles (on for 5 seconds and off for 5 seconds) on ice. Large proteins were removed by centrifugation at 10,000 xg for 15 minutes, and supernatant was transferred to new tube. The sample was freeze-dried using a vacuum centrifuge (Speedvac concentrator SVC200H, Savant, USA) and stored at - 80°C for further treatment.

### MALDI-TOF/TOF mass spectrometry analysis

The sample was prefractionated on a C18 column (BEH C18 column, Waters, Milford, MA, USA) using mobile phase at high pH (MilliQ and acetonitrile with ammonium hydroxide (20 mM), pH 10) and five fractions were made during the gradient (B: 5 to 90 % in 30 minutes) at a flow rate of 100 μL /min. Prefractionated samples were further separated on C18 column at low pH. The eluent was (A) water containing 0.5% FA and (B) 90% acetonitrile in 0.5% aqueous formic acid. The column was first washed and equilibrated with eluent A and 5% of eluent B. After loading the sample, a linear gradient from 5% B to 60% B in 60min at a flow rate of 100 μL /min was used as the mobile phase. Thirty fractions of 200 μL were collected from the beginning of the gradient using an automatic fraction collector. The resulting samples were dried in a Speed-vac and stored at −80°C until further analysis. Fractionated samples were resuspended in 1.5 μL water/ acetonitrile/FA (50/49.5/0.5 v/v). Subsequently they were transferred onto a MALDI target plate (Bruker Daltonics, Bremen, Germany) and mixed with 1.5 mL of a saturated solution of CHCA in 50% acetonitrile containing 0.5 % FA. After evaporation of the solvent, the MALDI target was introduced into the mass spectrometer ion source. Tandem mass spectrometry analysis were performed on an Ultraflex II instrument (Bruker Daltonics, Bremen, Germany) in positive ion, reflectron mode. The instrument was calibrated externally with a commercial peptide mixture (peptide calibration standard, Bruker Daltonics). All spectra were obtained using Flex Control software (Bruker Daltonics, Bremen, Germany). The plate was initially examined in MS1 mode and spectra were recorded within a mass range from m/z 500 to 4000. Subsequently, the peaks with S/N 10 were selected and used for the optimized LIFT method from the same target. All tandem mass spectra were processed by means of the FlexAnalysis software (Bruker Daltonics, Bremen, Germany), and m/z values and intensities of each peak were recorded in peak list files.

### Mass spectrometry data analysis

In this study, potential neuropeptide sequences are systematically extracted from proteins in order to reduce the search space in peptide-spectrum matching as follows. Amino acid sequences of proteins predicted from the genome of *Nematostella vectensis* were obtained from JGI (*Nematostella vectensis* version 1.0, all models) [20]. An in-house software module was developed enabling the prediction and extraction of potential mature neuropeptide sequences from the *Nematostella* proteins. First, the amino acid sequence of a protein is scanned for residues corresponding to input amino acid pattern (e.g. GK/R for C-terminal amidation) in a sliding window fashion. Secondly, an amino acid motif corresponding to an N-terminal structure (e.g. X(X)P) is searched at the N-terminal side from C-terminal motif found within a given amino acid length. In case multiple locations found to match N-terminal motif, all of them are considered to be potential N-termini. All combinations of input amino acid patterns are applied. All amino acid sequences that meet the criteria are cleaved *in silico* and written in a FASTA formatted file and used as target database for peptide-spectrum matching. For the present study, the following amino acid motifs were used to extract potential neuropeptide precursor protein sequences.

C-terminal motif

G[K/R]↓: glycine loss-amidation N-terminal motif

↓XP: proline from the second position from N-terminus

↓XXP: proline from the third position from N-terminus

↓Q: Glutamate for the formation of pyroglutamic acid

(↓ indicates the cleavage site)

The software was coded in C++ using Visual Studio 2008 (Microsoft) and Boost library. The software is available upon request.

All tandem mass spectra were searched against a database holding (A) the entire proteins and (B) the predicted neuropeptide sequences, using the Mascot search engine version 2.3 (Matrix science, London, UK). Peptide-spectrum matching were performed allowing variable modifications of glycine-loss amidation, pyroglutamic acid (Glu), and oxidation (Met), which are common post-translational modifications of neuropeptides. Searches against the predicted neuropeptide sequence dataset were performed with “No Cleavage” setting, in which cleavage of input sequences is not considered and precursor masses are matched to the theoretical masses of intact peptide sequences. Mass tolerance for precursors and fragment ions were set to 0.4 and 0.8 Da. In this study, peptides were first tentatively identified with a Mascot expect value less than 0.05 and then further confirmed by manual verification of the product ions assigned. Decoy search was conducted by Mascot decoy database function. Signal peptides of the precursors of identified peptides was examined using SignalP version 4.1 [21].

### Expression analysis

Partial ORF sequences for the identified peptides were obtained from the National Center for Biotechnology Information (NCBI) trace archive of the *Nematostella vectensis* (data generated by the Joint Genome Institute) and from *Stellabase* (http://cnidarians.bu.edu/stellabase/index.cgi). Gene-specific primers were designed based on the ORF sequences. The primer sequences are available upon request. PCR products for the peptide genes were subcloned into pGEM-T (Promega) and sequenced. Riboprobes were synthesized and purified as described previously [22]. *In situ* hybridization was performed as described previously [23] with following modifications: specimens were fixed with 4% paraformaldehyde/PBS + 0.1% Tween 20 (PBST) for 1 hr, washed with methanol for 3 times and stored at −20°C. Hybridization of 0.4-to 1.1-kb digoxygenin (DIG)-labeled antisense RNA probes was carried out using hybridization solution containing 1% SDS at 50-65°C for at least 22 hrs. For post-hybridization washes, specimens were washed by serial dilutions (75%, 50%, 25%) of hybridization solution with 2× SSC at 55°C. After DIG-labeled probe was visualized using BM purple (Roche), specimens were washed with PBST.

## Results

### Strategy

Systematic neuropeptide identification was performed for *Nematostella vectensis*, using a combination of neuropeptide prediction combined with tandem mass spectrometry. Fig 1 shows an overview of the used approach. Protein database of *Nematostella vectensis* were first processed with our newly developed neuropeptide prediction tool to extract potential neuropeptide sequences based on their structural hallmarks such as cleavage sites and N/C-terminal PTMs. Fragmentation spectra were acquired by means of LC-MALDI TOFTOF. Unlike conventional peptide-spectrum matching for endogenous peptides or enzymatic digests, we narrowed the search space and only used the sequences of *in silico* cleaved potential neuropeptides a target sequence dataset for peptide-spectrum matching. Typically in peptidomics, searches are often done considering any amino acid residue as potential cleavage sites. In the present method, the search is done directly against the mature forms of potential neuropeptide sequences in a manner similar to top-down analysis of protein.

**Fig 1.**
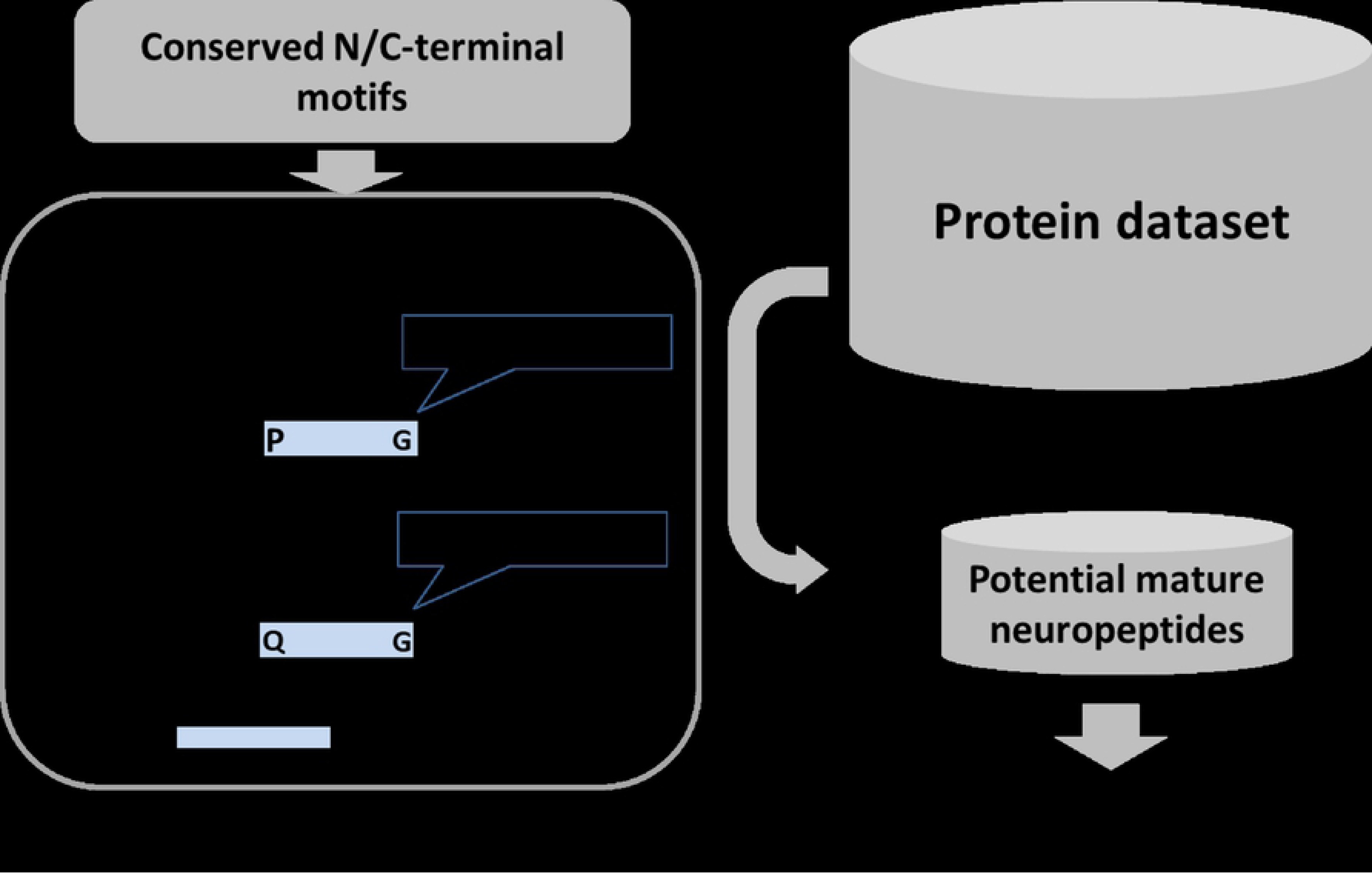
Scheme of the neuropeptide identification strategy using a combination of *in silico* cleaved neuropeptide predictions and peptide-spectrum matching. The N/C-terminal motifs are used to scan the protein database for the candidates of neuropeptides. The detected sequences are exported as a target database for top-down MS/MS search.

### Building prediction models

To find common structural hallmarks of neuropeptides in cnidarian species, we collected neuropeptide sequences from the UniProt sequence database and other publically available resources (supporting table 1)[24–33]known neuropeptides in cnidarian species, 52 % have N-terminal prolines and 54 % have N-terminal glutamines, which are used for the conversion to pyroglutamic acid. In total, 84 % of cnidarian neuropeptides have either proline or glutamine at N-termini, indicating these structures can be hallmarks to effectively extract potential neuropeptide sequences from protein sequences. All of the known cnidarian neuropeptides contained an amidation at their C-termini, which is a commonly observed PTM for neuropeptides in all species across the Animal Kingdom. As indicated by many studies, cleavages at dibasic residues ([KR] X{0,2,4,6}[KR]), which is the target site for the prohormone convertases, is commonly used to generate neuropeptides from precursor proteins.[1] This processing pattern can be found in peptide precursor proteins of many bilaterian species.[9] It should be noted, however, that in cnidarians dibasic residues are rarely used as cleavage sites within neuropeptide precursors (supplemental table 1). They do not show particularly conserved cleavage sites. Based on these observations, we selected N-terminal protective structures (Q or E for pyroglutamic acid formation or P at the second or third position from the N-termini) as well as C-terminal amidation (glycine and basic residues (K or R) as structural hallmarks for neuropeptide prediction. This is because these structures can be found in the vast majority of neuropeptides in cnidarian species and are not limited to particular neuropeptide families (supplemental table 1), thereby enabling comprehensive and untargeted neuropeptide identification. This information (types of amino acid residues and locations on peptides) was fed to our neuropeptide prediction tool as input, and protein database of *Nematostella* were scanned for potential neuropeptide sequences using the combination. Without any cleavage site specification (conventional peptidomics search), about 100,000 to 200,000 peptide sequences were found to fit the molecular masses of each observed peptide in the sample with the given mass tolerance in the *Nematostella* proteins. By comparison, in the newly constructed database that contained only *in silico* cleaved predicted neuropeptide sequences, only 400 to 800 peptides could be found. This efficiently reduced the size of the search space.

**Table 1.** Sequences of detected peptides and their Mascot E-value. E-values lower than the threshold (0.05) were indicated in bold. C-terminal amidations, oxidation, and N-terminal pyroglutamic acids are respectively indicated as “a”, “o” and “p”.

| Family name | Sequence | Mascot e-value |  |  |
| --- | --- | --- | --- | --- |
|  |  | predicted peptides | protein database | basic cleavage sites |
| HIRamide | APPLDLSGPAYFHIRa | **1.9E-02** | 4.2E+00 | nd |
|  | GPPYIDLTEPSFFHIRa | **8.8E-09** | **2.1E-06** | nd |
|  | NPPIDLGPAYFHIRa | **7.3E-07** | **1.6E-04** | **5.9E-07** |
|  | pQPPIDLSPAAYFHIRa | **1.1E-04** | **2.5E-02** | **7.7E-05** |
|  | pQPPLDLGPAYFHIRa | **5.0E-07** | **1.1E-04** | nd |
|  | pQPPYLDLGEPSFFHIRa | **4.5E-04** | 1.0E-01 | **3.6E-04** |
|  | pQPPYLDLTPAYFHIRa | **2.5E-04** | 5.5E-02 | nd |
|  | pQPPYLDLTPSYFHIRa | **5.4E-06** | **1.2E-03** | nd |
|  | pQPPMIDLSEPAFFHIRa | **1.2E-06** | **2.7E-04** | nd |
|  | pQQPPMIDLSEPAFFHIRa | **1.7E-03** | 3.5E-01 | nd |
| PRGamide | GPRGGRATEFGPRGa | **1.2E-04** | **2.8E-02** | nd |
|  | GPRGGREVNLEGPRGa | **6.7E-03** | 1.7E+00 | nd |
| QWamide | IPPQGFRFNQWa | **3.4E-02** | 8.2E+00 | nd |
|  | IPPQGLRFNQWa | **2.9E-03** | 6.7E-01 | nd |
|  | IPPQGLRFSQWa | **2.8E-03** | 6.4E-01 | nd |
| RFamide | MPEQDANPQTRFDa | **2.3E-05** | **5.3E-03** | nd |
|  | pQGRFGREDQGRFa | **3.2E-03** | 6.3E-01 | nd |
| RPamide | FPPGFHRPa | **2.0E-03** | 4.9E-01 | nd |
| RHamide | GPPMIKIPVRHa | **2.0E-03** | 5.0E-01 | nd |
|  | GPPMoIKIPVRHa | **7.7E-04** | 2.5E+01 | nd |

The tandem mass spectra were acquired from off-line LC-MALDI TOF-TOF analysis of a peptide extract of *Nematostella vectensis*. Peptide identification was carried out by using the Mascot peptide-spectrum matching tool, taking into account pyroglutamic acid formation, glycine-loss amidation and oxidation on M. The dataset of extracted potential neuropeptide sequences was used as target sequence dataset for peptide-spectrum matching, and the search was done in the top down fashion with “no-cleavage” setting. Twenty unique peptides were identified with a Mascot e-value < 0.05. Table 1 shows the identified peptide sequences. The amino acid sequences and the details of precursor protein coding genes can be found in supplemental table 2.

We compared the peptide identification performance of our approach with the conventional peptide-spectrum matching against the whole protein dataset in which every amino acid residue is considered as a potential cleavage site. As shown in table 1, the e-values of the search results using our predicted peptide dataset were significantly improved and our approach yielded more than twice as many peptide identifications. Fig 2 shows the overall distribution of the e-values of these peptides in the target and decoy searches. The e-values of the peptide hits in the protein database (Fig 2C) are distributed in the higher range and many of them show similar e-values as the decoy search result (Fig 2D), making the peptide sequence identification difficult. On the other hand, the e-values of the peptide hits using the reduced database of predicted candidate peptide sequences (Fig 2A) are distributed in a lower range, and show clear separation from the range of e-values of the decoy search result (Fig 2B).

**Fig 2.**
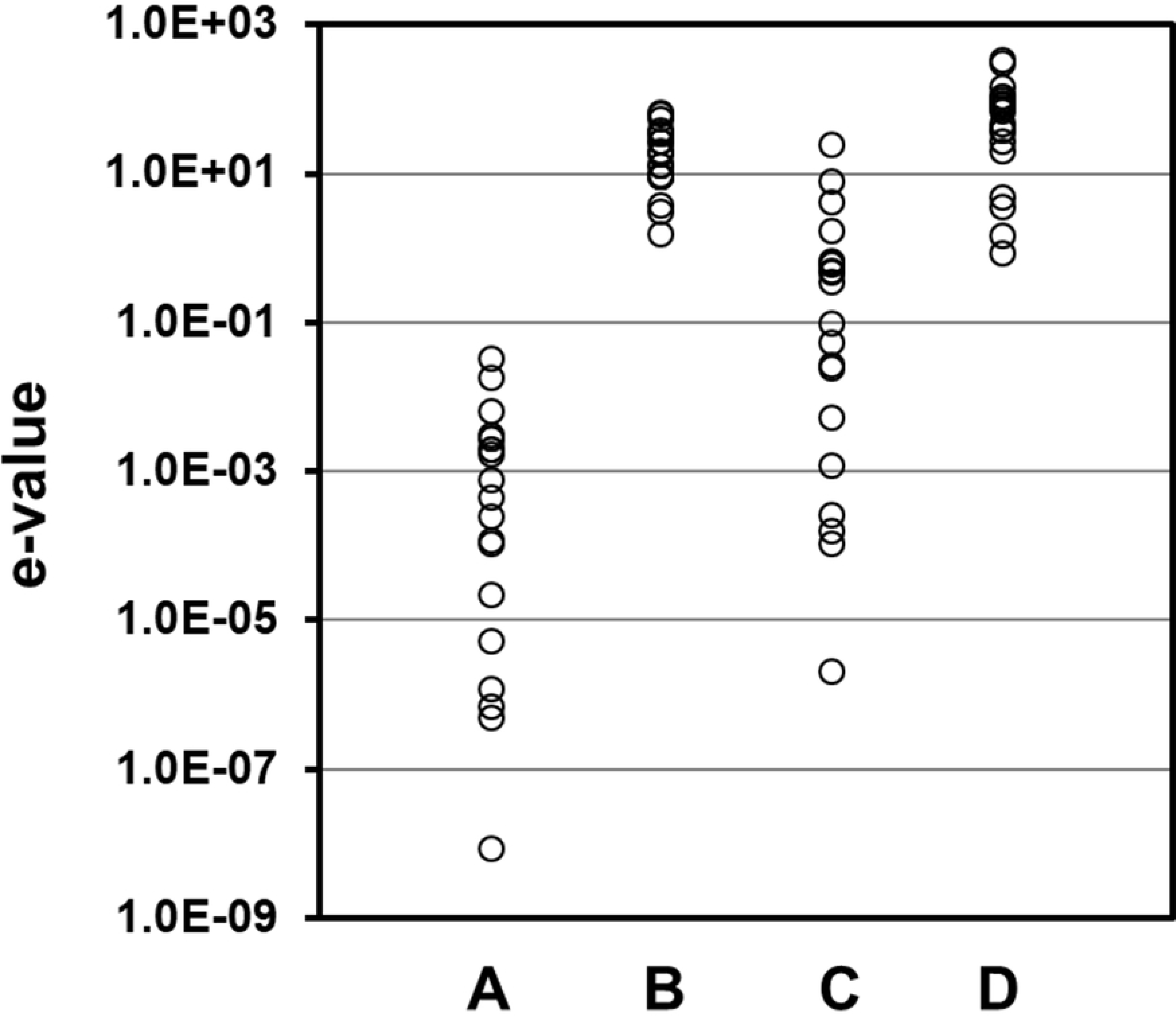
The distribution of e-values of the hits found in the datasets. Each circle indicates the e-value of the top-scoring hit in the dataset. A: Search against the predicted neuropeptide dataset. B: decoy of the predicted neuropeptides dataset. C: the whole protein models. D: decoy of the whole protein models.

In addition, to demonstrate the difference from the method in which only basic cleavage sites are used to narrow search space [34], we also scanned the protein database using the cleavage rule of prohormone convertases ([KR] X{0,2,4,6}[KR]). Among the peptides detected by using the structural hallmarks of neuropeptides, only 3 were identified (Table 1). This indicates that, narrowing down search space with a simple cleavage rule failed to predict neuropeptide sequences.. The peptide sequences and their precursors acquired in this study indicate that typical prohormone convertase cleavage sites (K/R) are not commonly utilized for the neuropeptide precursor processing in *Nematostella*, which agree with the neuropeptides found in other cnidarian species[25,30,35–37].

### Novel neuropeptides in *Nematostella vectensis*

Most of the peptides identified in the present study have not been reported yet, and their precursor protein-coding genes were not annotated as neuropeptide precursor, except for the RFamide precursor. Fig 3 shows the overview of the precursors that contain identified peptides. Overall, these precursors contain multiple copies of the mature peptide sequences sharing the same C-terminal structures. It should be noted that the precursors also contained multiple other potential neuropeptide sequences which contains same structural hallmarks used to the *in silico* cleavage, but not observed in this study. It is unclear whether those peptides are not produced or because of the sensitivity of mass spectrometry analysis is insufficient to detect those peptides.

**Fig 3.**
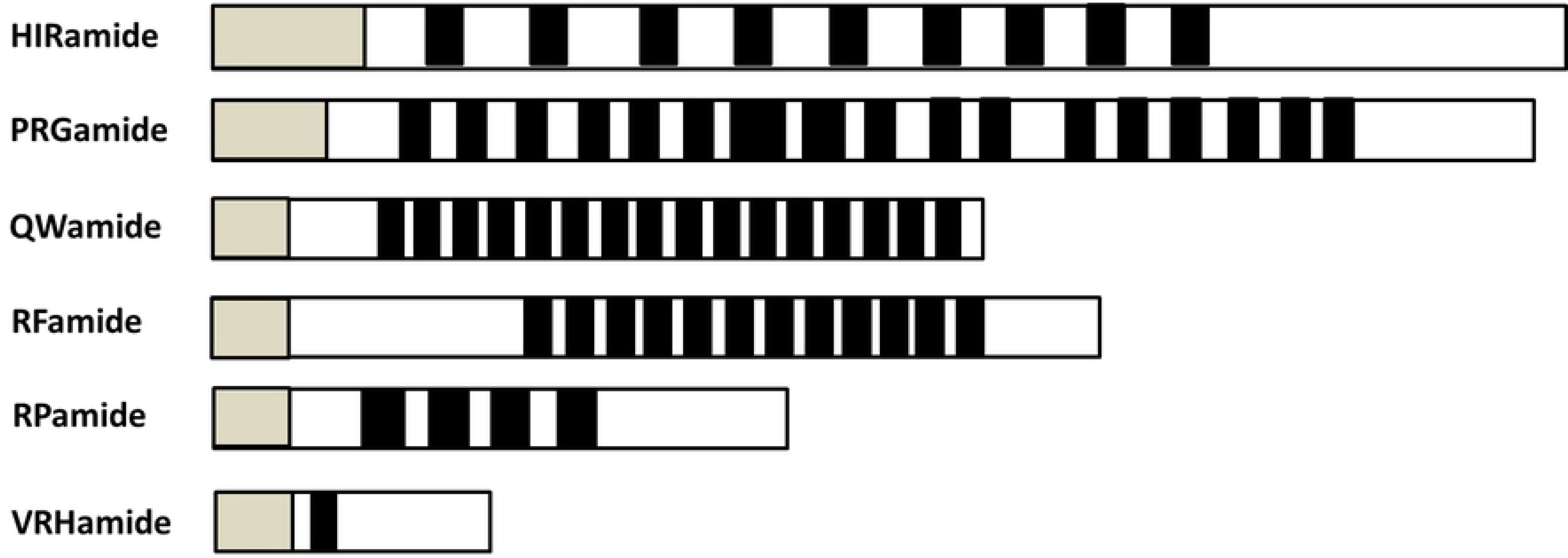
Overview the structures of neuropeptides precursor proteins detected in this study. Gray indicates the signal peptide. Black indicates the locations of the detected and potential mature neuropeptides.

The HIR peptide precursor contains 9 structurally related peptide sequences and a typical signal peptide sequence at the N-terminus. They share a high degree of structural similarity at the C-terminus with Arthropoda Tachykinin peptides (Fig 4A). Especially, *Locusta*-tachykinin-1[38] displays strong similarity to the *Nematostella* HIR peptides. Tachykinin and related peptides are well studied neuropeptides conserved in both invertebrates and vertebrates. Based on the C-terminal motifs, Tachykinin and related peptides can be further classified into two subgroups: the “-FXGLMa” and “-GFXGXRa” subfamilies. *Nematostella* HIR peptides show similarity to the “GFXGXRa” subfamily; especially the presence of phenylalanine or tyrosine residues, and C-terminal arginine is conserved in *Nematostella*. Arthropod and Nematostella share the PXXFYXXRamide motif.

**Fig 4.**
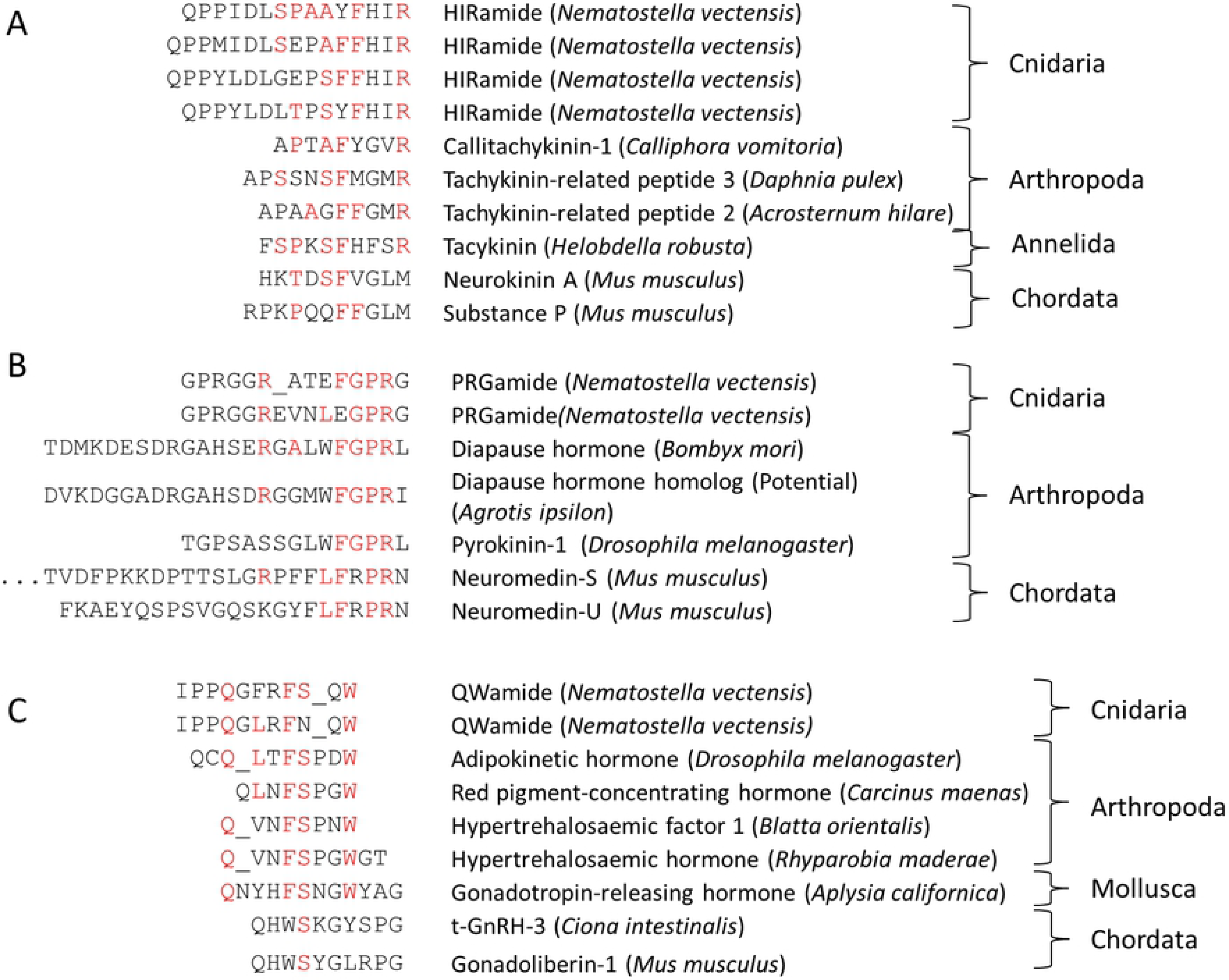
Structural similarities of identified neuropeptides in *Nematostella vectensis* and other species. A: HIRamides, Tachykinin related peptides[16,39–43]. B: PRGamides and PRXamide related peptides[44–48]. C: QWamides, AKH and Hypertrehalosaemic hormones [49–54]. Conserved amino acid residues are shown in red.

RFamides, a neuropeptide family widespread among bilaterian organisms and also previously reported in cnidarians including sea anemone, were also detected in this study. In this study an additional member of RFamide MPEQDANPQTRFDa was identified in the precursor protein of RFamide. This new member has an additional D at C-terminus, although all the other members are showing the RFamide motif. Two peptides, GPRGGRATEFGPRGamide and GPRGGREVNLEGPRG, share the C-terminal motif GPRGamide. Their peptide precursor contains a large number of peptide copies all sharing this motif. The peptides share sequence similarities with protostomian pyrokinins that are characterized by the C-terminal sequence FXPRLamide [55,56]. Pyrokinins are involved in the stimulation of gut motility, the production and release of sex pheromones, diapause, pupariation and feeding. (Fig 4B) [57] [58].

Three peptides derived from a single precursor protein display the C-terminal sequence motif FSQWamide or FNQWamide (Fig 4C). This motif aligns with the motif that typifies the bilaterian AKH/GnRH peptide family which is known to regulate energy metabolism and reproduction [59]..

In addition to the peptides described above, we discovered two groups of peptides (RPamide and RHamide) originated from two precursor proteins. We could not find any homologies to known neuropeptide families (table 1).

### Expression patterns of HIRamide, PRGamide, VRHamide neuropeptides

Our whole mounts in situ hybridization (WISH) analyses have confirmed that the peptide genes encoding the identified peptides (HIRamide, PRGamide, VRHamide) are exclusively expressed in neurons. As shown in fig 5, the expression of all peptide genes that could be detected by WISH was observed in specific cells housed mainly in the endodermal layer at the juvenile polyp stages. Careful observations at the cell morphology demonstrated that the cells exhibiting the expression of the genes shows a round-shaped cell body with neurite-like processes (Fig 5, lower panel). This indicates that these detected peptides are in fact expressed in neurons, and are neuropeptides. WISH experiments demonstrated also that the HIRamide and PRGamide neuropeptides are strongly expressed around the mouth opening (Fig 5a). This finding indicates that neuronal subsets expressing these neuropeptides develop at the oral side to form region-specific neural network. The tissue around mouth of cnidarian polyps have been shown to express a bunch of neuronal markers including RFamide, which is also evolutionarily conserved among metazoans (Fig 5, upper panel) [19], and to develop a regionalized nervous system which is known as the oral nervous system (*Nematostella*) or the nerve ring (*Hydrozoa*) [60–62]. Expression pattern of VRHamide encoding gene showed a sharp contrast to that of HIRamide, PRGamide and RFamide genes. The expression level of VRHamide gene at the oral tissue was under detection limit. Instead, it was strongly and exclusively detected in neurons located at the most distal region of the tentacle endoderm (Fig 5, upper panel). This unexpected and interesting expression pattern suggests the specific function of the VRHamide for development and/or contractile response of the tentacles (see Discussion).

**Fig 5.**
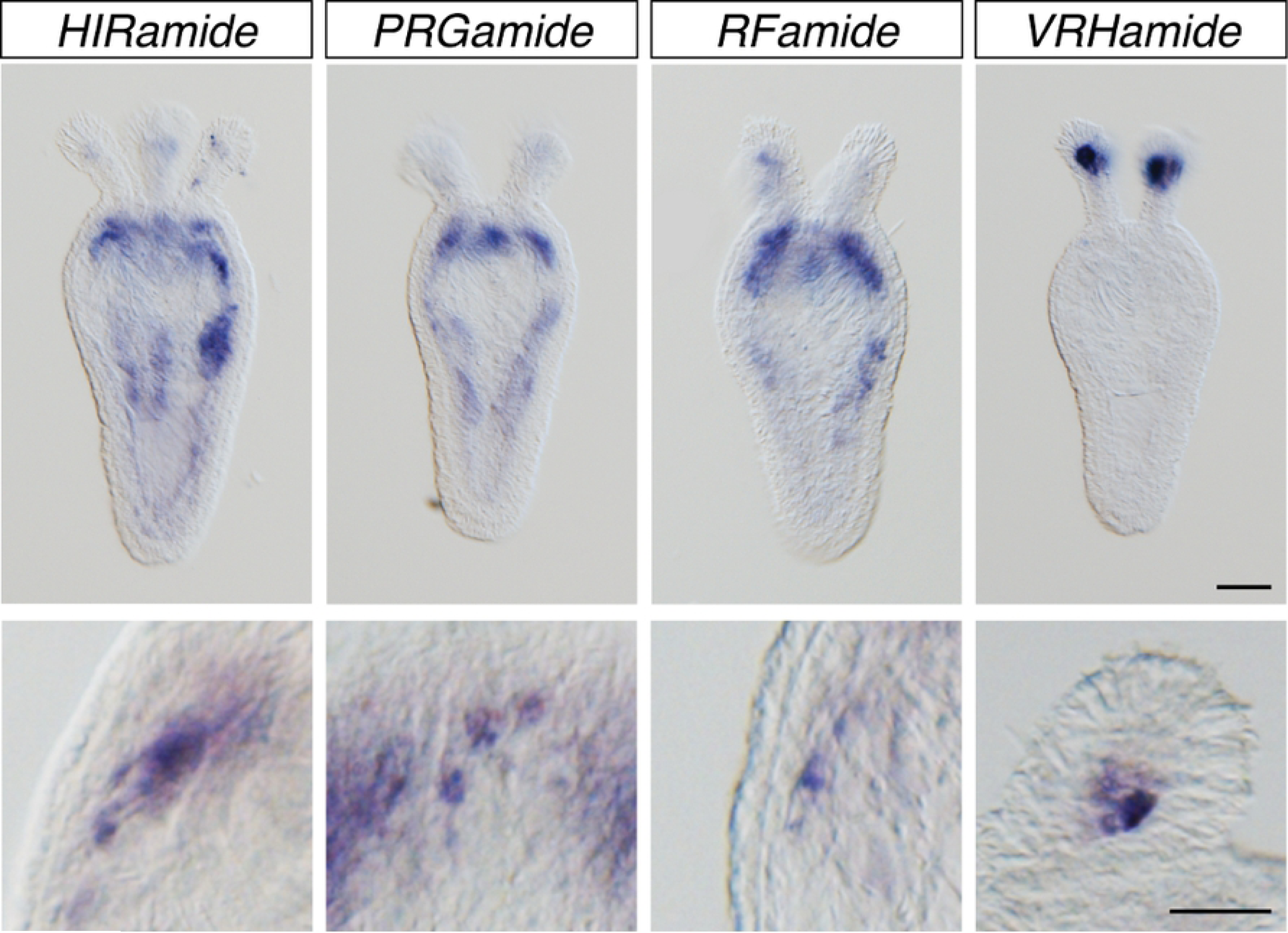
WISH staining of juvenile polyps of *Nematostella vectensis* (8 days post fertilization) showing localized expressions of *HIRamide, PRGamide RFamide* and *VRHamide* genes at low magnification (upper panels) and high magnification (lower panels). Scale bars, 100 μm (upper) and 50 μm (lower).

## Discussion

We studied the peptidome in *Nematostella vectensis*, which belongs to the Cnidaria that have one of the most basal nervous systems in the Animal Kingdom. Recent advances in mass spectrometry technologies have greatly enhanced the identification of neuropeptidomes from biological matrices [12,63–65]. However, an effective informatics solution for neuropeptide identification is still missing and leaving the interpretation of tandem mass spectral data still difficult. We overcame this difficulty by performing peptide spectrum matching against a database comprised of *in silico* cleaved putative predicted neuropeptide sequences. We successfully identified 20 peptides encoded in six neuropeptide precursor genes from *Nematostella* and confirmed experimentally that at least four precursor genes are expressed in neurons exhibiting distinct neurophysiological activities. Our approach can be easily incorporated with recent advanced technologies in peptidomics, and such a combination will greatly improve the analysis of the neuropeptidome in a broad range of biological materials.

### Strategies for the improvement of peptide identification

Peptide-spectrum matching has become a main tool for endogenous peptide identification. Unlike classical methods such as Edman degradation, this approach takes advantage of the high sensitivity of modern mass spectrometry instruments and high throughput peptide search engines and therefore has enabled comprehensive neuropeptide characterization. One of the main difficulties in peptidomics is that the peptide-spectrum matching needs to be done with the “no enzyme” setting, which results in an enormous search space. Consequently the peptide-spectrum matching result is regularly worse as compared to a peptide search in a proteomics setting.

The peptide characterization strategy employed in this study is a combination of predicting potential neuropeptide sequences and peptide-spectrum matching. This approach is effective in narrowing the search space and thereby significantly increases the sensitivity of peptide identification. Fälth *et al.* employed a similar approach to narrow search space in order to improve peptide-spectrum matching [34]. They used the most common cleavage site rule to extract potential neuropeptide sequences from a large protein database, and significantly improved peptide identification rate. Southey *et al.* developed a dedicated tool to predict the basic cleavage sites in neuropeptide precursor genes.[66] The prediction engine, NEUROPRED, needs to be trained on the information of known neuropeptides and their precursors; hence a large number of identified peptides and the information of processing sites are required. Compared to the prediction with simple motif matching, the prediction by NEUROPRED using the logistic regression modelling is more reliable, but on the other hand rather strict and potentially discards many forms of cleaved peptides especially if the large amount of training data is not available. These approaches are effective for the prediction of neuropeptides in well-studied animal phyla. On the other hand, it’s difficult to predict peptides from less-well studied animal models such as cnidarians. Unlike bilaterian animals, non-basic cleavage sites are commonly observed in cnidarian neuropeptide precursors [25,27,31,33,35–37,67]. In fact, most of the peptides identified in the present study are generated via cleavage at non-basic residues and unconventional sites. Therefore, the approach we took in the present study, i.e. the prediction of putative neuropeptide sequences based on structural hallmarks such as protective N-terminal pGlu or X(X)P and C-terminal amidation is more comprehensive and effective to preserve as many potential neuropeptide sequences as possible as the target database for peptide-spectrum matching. Evidently, the approach has also a drawback as it only allows identification of all neuropeptides that display these specific features. Non-amidated peptides or peptides that are not N-terminally protected are not in the database. At this moment, there are no means other than homology-based searches, or Neuropred prediction tools to search for the latter peptides, and thus far these tools are ineffective for cnidarian neuropeptides because of aforementioned reasons.

### Neuropeptide evolution

Many pioneer studies in the late 80ties and early 90ties already pointed to homologies between protostomian and deuterostomian neuropeptides based on neuropeptide sequence identifications in vertebrate, insect, molluscan and cnidarian species [18,30,68–72]. However, until today knowledge on signalling molecules, including neuropeptides, in eumetazoan animals is scarce. A previous study showed a highly conserved set of genes in the *Nematostella vectensis* genome, including molecules involved in neurotransmission [19,73]. The genome displays a number of receptor genes such as G-protein coupled receptors showing high degrees of similarities to known neuropeptide receptors. It is therefore highly likely that this primitive animal contains a variety of signalling peptides. Anctil *et al.* has reported potential neuropeptide coding genes by using homology searching [74]. Although those predicted precursor proteins contain short repeats of neuropeptide-like motifs, it is difficult to prove their identities as these neuropeptide precursors were predicted solely based on their protein sequence without any experimental evidence. In fact some of the predicted peptide precursor proteins in this study show a significant degree of structural similarity to non-neuropeptide precursor proteins, making it difficult to decide whether they are true neuropeptide precursors. In the present study we were unable to identify any of those predicted peptides, either by conventional “no-enzyme” searches nor by searches against the predicted neuropeptides. Homology-based searches based on short neuropeptide sequences is thus still difficult and, more importantly, the homology-based prediction of neuropeptide precursor genes does not provide evidence for the occurrence of predicted neuropeptides *in vivo*. Therefore, peptide identification approaches that are based on conventional biochemical purification methods and powered by upcoming bioinformatic technologies are necessary to empirically prove the predicted peptides and will be important to identify not only evolutionary conserved peptides but also evolutionarily derived or species-specific peptides that correspond to the diverged physiological traits.

Our data have also important implications on our insight into the evolution of neuropeptides. Since the neuropeptide repertoire in ancient eumetazoans is largely unknown, the origin of most of neuropeptides found in bilaterian animals remains difficult to reveal. Jekely (2013) performed a similarity-based clustering analysis of genes encoding neuropeptides and neuropeptide GPCRs across metazoan phyla and concluded that the last common ancestor of eumetazoans had various small amidated peptides including RFamide, RYamide, and Wamide) [75]. Ancestral urbilaterian neuropeptide-receptor families include GnRH, vasopressin, GnIH/SIFamide, CRF/diuretic hormone, calcitonin/DH31, NPY/NPF, neuromedin-U/pyrokinin, CCK/sulfakinin, galanin/allatostatin-A, and orexin/allatotropin [75,76]. It was suggested that these neuropeptide families may have originated concomitantly with the origin of a complex bilaterian body plan having a through gut with novel controls for food intake and digestion, excretory and circulatory systems, light-controlled reproduction, a centralized nervous system, complex reproductive behavior, and learning.

The present study indicates that the extent of the conservation of some neuropeptide families is even deeper as proposed in aforementioned studies, as we were able to identify neuropeptides of the GnrH/AKH, tachykinin and neuromedin-U/pyrokinin families in a cnidarian Nematostella, suggesting that these neuropeptide families were already present in the common ancestors of all eumetazoan species. Because *Nematostella vectensis* is a member of the Anthozoa, the most basal cnidarian group, the neuropeptides identified in the present study might constitute some of the most ancient neuropeptide in animal evolution. At least the tachykinin, GnRh, neuromedin-U and RFamide neuropeptides may have originated concomitantly with the formation of a nervous system. All these neuropeptides play a variety of roles in animal physiology, including feeding and metabolism, locomotion, reproduction, homeostasis, biorhythms and associated behaviors. We anticipate that the discovery of the receptors, downstream targets, and functions of cnidarian representatives of ancient eumetazoan neuropeptide families will provide important new insights into the evolution of neuropeptide function in the Animal Kingdom

## Acknowledgement

This work was supported by the agency for Innovation by Science and Technology (IWT Flanders), the Industrial Research Fund of the KU Leuven, and the FWO (Flemish Research Foundation).

## Supporting Information

**S1 Table.** The structures of the cnidarian neuropeptides.

**S2 Table.** Details of the neuropeptide precursor genes.

